# Genome-wide association study reveals genetic link between diarrhea-associated *Entamoeba histolytica* infection and inflammatory bowel disease

**DOI:** 10.1101/137448

**Authors:** Genevieve L Wojcik, Chelsea Marie, Mayuresh M Abhyankar, Nobuya Yoshida, Koji Watanabe, Josyf Mychaleckyj, Beth D Kirkpatrick, Stephen S Rich, Patrick Concannon, Rashidul Haque, George C. Tsokos, William A Petri, Priya Duggal

## Abstract

Diarrhea is the second leading cause of death for children globally, causing 760,000 deaths each year in children under the age of 5. Amoebic dysentery contributes significantly to this burden, especially in developing countries. We hypothesize that genetic variation contributes to susceptibility to diarrhea-associated *Entamoeba histolytica* infection in Bangladeshi infants; thus, we conducted a genome-wide association study (GWAS) in two independent birth cohorts of diarrhea-associated *E. histolytica* infection. Cases were defined as children with at least one diarrheal episode positive for *E. histolytica* through either PCR or ELISA within the first year of life. Controls were children without any episodes positive for *E. histolytica* in the same time frame. Meta-analyses under a fixed-effects inverse variance weighting model identified variants in two neighboring genes on chromosome 10: *CUL2* (cullin 2) and *CREM* (cAMP responsive element modulator) associated with *E. histolytica* infection, with SNP rs58000832 achieving genome-wide significance (P_meta_=4.2x10^−10^). Each additional risk allele (an intergenic insertion between *CREM* and *CCNY*) of rs58000832 conferred 2.5 increased odds of a diarrhea-associated *E. histolytica* infection. The most associated SNP within a gene was in an intron of *CREM* (rs58468685, P_meta_=2.3x10^−9^), which with *CUL2*, has been implicated as a susceptibility locus for Inflammatory Bowel Disease (IBD) and Crohn’s Disease. Gene expression resources suggest these loci are related to the higher expression of *CREM*, but not *CUL2*. Increased *CREM* expression is also observed in early *E. histolytica* infection. Further, *CREM*^-/-^ mice were more susceptible to *E. histolytica* amebic colitis. These genetic associations reinforce the pathological similarities observed in gut inflammation between *E. histolytica* infection and IBD.

## Introduction

Despite drastic reductions in childhood mortality from 1990 to 2015, the Millennium Development Goal 4 failed to reach the target goal of 2/3rd reduction in under-5 mortality rate. ^1^. One of the leading causes of childhood mortality is diarrheal disease, leading to over half a million deaths annually.^2^ The burden of diarrhea-related mortality and morbidity are disproportionately higher in developing countries. One endemic etiology of diarrheal disease in the developing world is amebiasis, an infection by the protozoan parasite *Entamoeba histolytica*.^3^ Observational studies have shown that preschool age children with a history of *E. histolytica*-associated diarrheal illness were more likely to be malnourished and stunted.^4,5^ While the majority of infections by *E. histolytica* are asymptomatic ^6^, the 10% that develop disease can exhibit acute diarrhea, dysentery, amoebic colitis and amoebic liver abscess.^7^

A major risk factor for amebiasis include poor sanitation and hygiene, as transmission occurs via the ingestion of amoebic cysts that are found in contaminated food and water.^8^ Within a group of 147 infants followed for the first year of life in an urban slum of Dhaka, Bangladesh, 10.9% of children had at least one diarrheal episode positive for *E. histolytica*. ^9^ Children were more likely to be infected with *E. histolytica* if they were born malnourished. These findings are consistent with prior evidence in preschool age children, which showed children with *E. histolytica*-associated diarrheal illness were more likely to be malnourished and stunted. ^5^ By the end of three years of follow-up (from ages 2 to 5), 17% of children had at least one diarrheal episode positive for *E. histolytica*. Within a homogenous environment with presumed uniform exposure to the pathogen, it is not well understood why only a subset of individuals exposed exhibit infection, and subsequently why symptomatic disease develops only in a subset of those individuals.

One possible explanation for the observed heterogeneity in infection rates could be differences in host genetic susceptibility to infection and disease.^10^ In a study of preschool age children in Dhaka, Bangladesh, Duggal et al identified a polymorphism in the leptin receptor, rs1137101 resulting in Q223R, was associated with amebiasis when compared to children without this polymorphism.^11^ Later work elucidated the mechanism of action of this allele; glutamine at this position led to a decrease in STAT-3-dependent gene expression, which in turn led to an increase in host cell apoptosis during *E. histolytica* infection.^12,13^ An additional association was found between human leukocyte antigen (HLA) class II alleles and *E. histolytica* infection in Bangladeshi children, specifically DQB1*0601 and the haplotype containing the DQB1*0601/DRB1*1501 heterozygote. ^14^ To comprehensively identify loci conferring risk for diarrhea-associated *Entamoeba histolytica* infection, we conducted a genome-wide association study by implementing a meta-analysis of 2 birth cohorts of children in Dhaka, Bangladesh: PROVIDE (Performance of Rotavirus and Oral Polio Vaccines in Developing Countries) Study^15^ and the Dhaka Birth Cohort^9^ (DBC).

## Results

### Outcome information

Within the well-characterized DBC and PROVIDE studies, children were followed twice weekly for a possible diarrheal episode. When a mother reported diarrhea in her child, a fecal sample was tested for amebiasis. Within DBC, 374 children were reported to have diarrhea; 65 of these children had at least one diarrheal episode positive for *E. histolytica* within the first year of life. A total of 309 controls had a stool sample collected within the first year of life (diarrheal or normal monthly) and they were not positive for *E. histolytica*. Within the PROVIDE study, 112 children had at least one diarrheal episode positive for *E. histolytica*, while 323 did not have any *E. histolytica* positive samples within the first year of life. **(Table 1)** Diarrhea-associated *E. histolytica* infection had no association (*P*> 0.05) with height-for-age Z-score (HAZ) at one year of age, the number of days that the child was exclusively breastfed, or sex. **(Table 2)** Results were consistent between the DBC and PROVIDE (*P*_*het*_ > 0.05).

**Table 1:** Outcome distribution of cases and controls within the two cohorts.

| Study | Case | Control | Total |
| --- | --- | --- | --- |
| Dhaka Birth Cohort | 65 | 309 | 374 |
| PROVIDE | 112 | 323 | 435 |
| **Total** | **177** | **632** | **809** |

**Table 2:** Association between possible covariates and outcome. (HAZ= Height-for-age Z-score)

| Covariate | DBC |  |  | PROVIDE |  |  | P<sub>het</sub> |
| --- | --- | --- | --- | --- | --- | --- | --- |
|  | Mean controls | Mean cases | P | Mean controls | Mean cases | P |  |
| HAZ<sub>12 months</sub> | -1.978 | -2.04 | 0.803 | -1.446 | -1.498 | 0.649 | 0.775 |
| Number of days of exclusive breastfeeding | 122.79 | 128.92 | 0.510 | 125.7 | 122.7 | 0.644 | 0.442 |
| Sex | 54.49% | 55.38% | 0.919 | 54.49% | 51.79% | 0.621 | 0.671 |

### GWAS meta-analysis identifies significant association in *CREM-CUL2* region

A total of 9.8 million SNPs with a pooled (across DBC and PROVIDE) minor allele frequency (MAF) greater than 1% were analyzed for association with diarrhea-associated *Entamoeba histolytica* infection within the first year of life. A genome-wide association analysis was performed separately for each study (DBC and PROVIDE) using an unadjusted logistic regression assuming an additive model, followed by a fixed effects meta-analysis. **(Figure 1)** The top genetic association (rs58000832, P_meta_=4.2x10^−10^, MAF=23%) was identified on chromosome 10 in a region covering three genes: *CUL2* (cullin 2), *CREM* (cAMP responsive element modulator), and *CCNY* (cyclin Y) **(Figure 2).** Individuals with at least one copy of the CA insertion at rs58000832, an intergenic insertion between *CREM* and *CCNY*, had 2.5 times increased odds of diarrhea-associated infection within the first year of life when compared to individuals with no copies of this insertion. The most associated SNP located in a gene (rs58468685, P_meta_=2.3x10^−9^) was in an intron of *CREM*. Within this region of association, there was a large linkage disequilibrium block underlies *CREM* and *CUL2*, flanked by two recombination peaks. **(Figure 2)**

**Figure 1:**
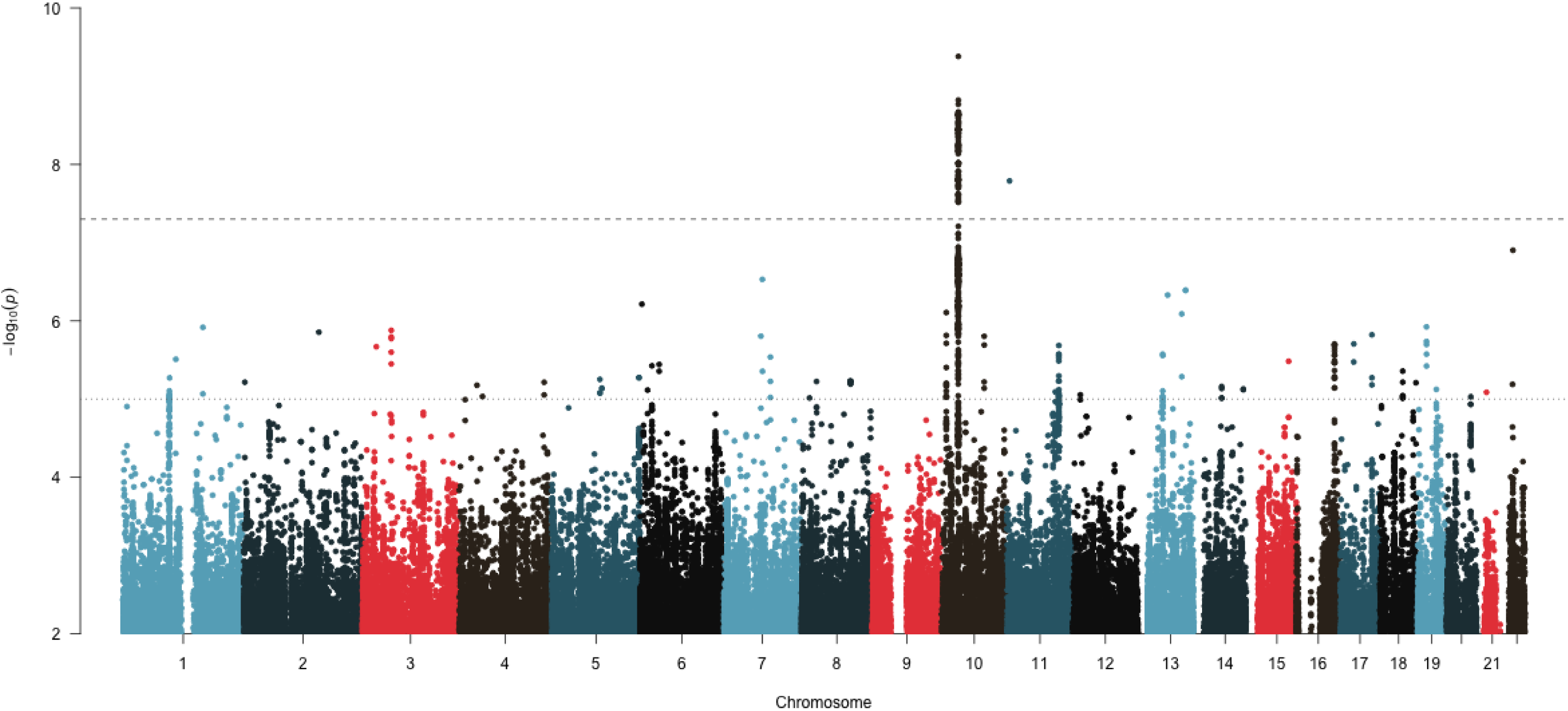
Manhattan plot of diarrhea-associated E. *histolytica* infection within the first year of life. The genome-wide significance threshold is denoted with the dashed line at 5×10^−8^, and suggestive significance threshold with the dotted line at 5×10^−5^.

**Figure 2:**
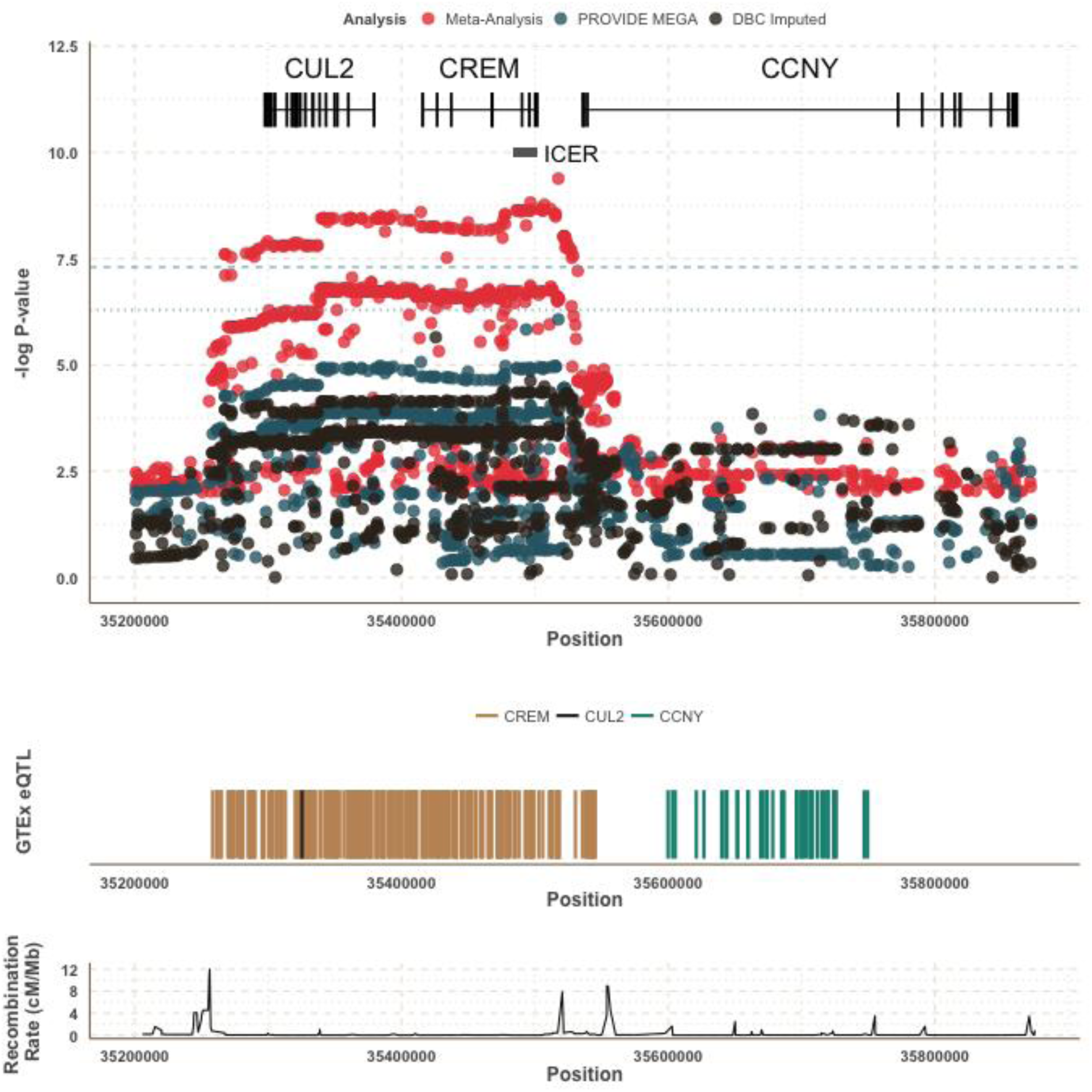
Zoomed-in Results of *CUL2-CREM-CCNY* Region and GTEx results of known expression quantitative trait loci (eQTL).

### *CREM/CUL2* haplotypes enriched for South Asian populations

There were 11 distinct haplotypes in the *CREM*/*CUL2* region within PROVIDE using the 26 SNPs of interest on chromosome 10 (physical positions: 35,273,439-35,513,323). The most associated haplotype (*P*=1.07x10^−4^), haplotype A, conferred 2.05 times increased odds of diarrhea-associated *E. histolytica* infection with each copy of the haplotype. This single haplotype encompasses both *CREM* and *CUL2*. When the haplotype was split into the two genic regions (*CREM* and *CUL2,* respectively), there was little difference in association results, likely due to the linkage disequilibrium across the region. **(Supplementary Table 1)** Each additional copy of the most associated *CUL2* haplotype conferred 2.2 times increased odds of diarrhea-associated E. *histolytica* infection (P=2.08x10^−5^), while each additional copy of the most associated *CREM* haplotype conferred 2.14 times increased odds of infection (*P*=3.67x10^−5^).

Haplotype A, the most associated haplotype across both *CREM* and *CUL2,* was examined in 5 main continental groups: Africa, the Americas, East Asia, South Asia, and Europe from the 1000 Genomes Project.^16^ Using the same SNPs, there were 23 distinct haplotypes seen at least 10 times within the 2,432 unrelated individuals. The risk haplotype from the PROVIDE analysis was found at the highest frequency in South Asian (6.1%) and European (4.4%) populations. East Asian (2.6%) and African (3.8%) populations had lower frequencies of the risk haplotype. A closely related haplotype (only one nucleotide difference from the risk haplotype) was also enriched within South Asian and European populations (12.8% and 12.1%, respectively), with lower frequencies (<5%) in both East Asian and African populations. **(Supplementary Table 2)** These haplotype results suggest that the risk haplotype is enriched within South Asian and European populations. However, there was no evidence supporting either directional or balancing selection. **(Supplementary Figures 1-2)**

### eQTL analysis reveals a role for *CREM*

Within the associated region of *CUL2*-*CREM*-*CCNY* were known cis-eQTLs (expression quantitative trait loci) identified by the Genotype-Tissue Expression (GTEx) Consortium.^17^ While known eQTLs in this region included representation of all three genes (*CREM*, *CUL2*, *CCNY*), the only sites that overlapped with our genetic associations from meta-analysis were with *CREM*. **(Figure 2)** Of the 490 cis-eQTLs for *CREM* in this region, 167 were significantly associated within the meta-analysis (*P*<5x10^−7^). In contrast, none of the *CUL2* (n=2) or *CCNY* (n=80) eQTLs that overlapped with an associated GWAS SNP (*P*<0.001) reached significance. This suggests that the GWAS associated SNPs may be related to *CREM* expression, and not *CUL2* expression The majority of *CUL2* eQTLs were within *CCNY* and, therefore, outside the area of the association. All *CREM* eQTLs were in a single cell type: EBV-transformed lymphocytes. Of the SNPs with overlap between the GWAS and GTEx eQTLs, the strongest effect was at rs12248333 (intronic within *CUL2*). The minor allele (G) has MAF=0.35, and the presence of each allele is was associated with increased odds of diarrhea-associated *E. histolytica* (*P*=8.8x10^−8^) and decreased expression of *CREM* (*P*=3.8x10^−6^). This inverse relationship was consistent for all overlapping eQTLs.

### *E. histolytica* activates cAMP signal transduction and induction of CRE-driven promoter elements *via* CREM

Previous work has shown that amebic lysates induce significant cAMP elevation in rat colonic mucosa^18^ and that secretory products of the parasite increase cAMP in leukocytes^19^. We hypothesized that CREM is activated by parasite-induced cAMP signal transduction. To investigate this hypothesis, we analyzed transcriptional activation of conserved cAMP-response elements (CRE) by *E. histolytica* in intestinal epithelial cells expressing a CRE-luciferase reporter. *E. histolytica* induced robust CRE activation after 1 hour (39.55 +/- 3.07 fold induction), which reached ∼700 fold after 4.5 hours (709.2+/- 53.94 fold induction). CRE-activation and repression by *E. histolytica* and the positive control forskolin displayed similar kinetics. **(Figure 3A)** To determine the contribution of CREM to *E. histolytica* CRE-activation, we measured CRE-activity in HCT116 cells silenced for CREM. Cells silenced for CREM had reduced CRE-activation in response to *E. histolytica* secreted products (24.1% of control, *P*=0.032) and forskolin (50.3% of control, P=0.009). **(Figure 3B).** These data suggest HCT116 intestinal epithelial cells induce a transcriptional response to *E. histolytica* via cAMP signal transduction activation of CREM.

**Figure 3:**
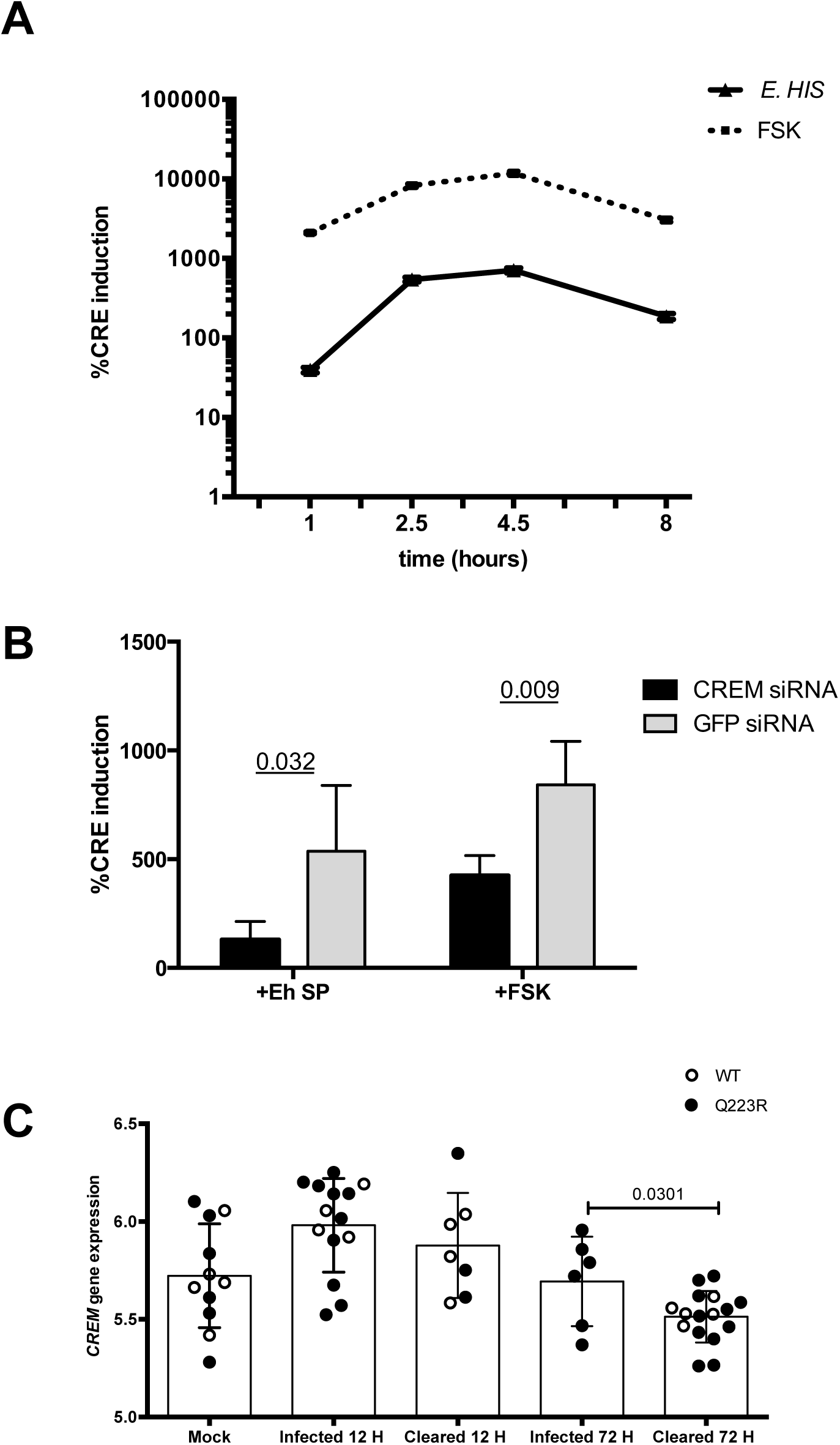
CREM is activated by *E. histolytica* secreted factors and CREM expression correlates with *E. histolytica* clearance in mice. **(A) *E. histolytica* activates CRE-driven gene transcription in intestinal epithelial cells.** HCT116 intestinal epithelial cells transfected with a cAMP-Response-Element (CRE) driven reporter exposed to *E. histolytica* trophozoites (*+E.HIS*) or 10 micromolar forskolin (+FSK), positive control for CRE-activation **(B) Silencing CREM decreases CRE reporter induction by soluble *E. histolytica*** indicating that CREM is the major transcriptional regulator acting at CREs in response to amebic secreted products. CREM was silenced by 96% compared to GFP controls by qPCR (data not shown/supplemental?) **(C) *CREM* expression is induced in early infection in mice.** Mice with a wild-type humanized leptin receptor gene (WT) or susceptible leptin receptor (Q223R) were infected with *E. histolytica* trophozites by intracecal injection and sacrificed at 12h or 72 h post infection. *E. histolytica* infection was measured by culture of cecal contents*. CREM* gene expression data from microarray analysis is shown. (Data from Mackey-Lawrence et al 2013).

### Expression of *CREM* during amebiasis

To elucidate how *CREM* impacts susceptibly *in vivo* we compared *CREM* expression in wild-type mice that are naturally resistant to *E. histolytica* and mice with increased susceptibility due to a single amino acid substitution (Q223R) in the leptin receptor. Mice were sacrificed at 12 and 72 hours post-infection. 50% (4/8) of wildtype mice were infected 12 hours post infection and none were infected 72 hours post infection (0/5 infected); in contrast, 77% (10/13) of the susceptible Q223R mice were infected 12 hours post-infection and 35% (6/17) remained infected 72 hours post-infection.^12^ *CREM* expression was increased in both susceptible and wildtype resistant mice 12 hours post-infection regardless of parasite clearance. After 72 hours, *CREM* expression was significantly lower in mice that had cleared the infection (both wildtype and susceptible) relative to mice that were still infected (Q223R only) (-0.18 +/-0.08, *P*=0.031) **(Figure 3C).**

### CREM knockout mice more susceptible to amebic colitis

To further determine the role of CREM in vivo, we compared the rates of amebic colitis between CREM knockout (C57BL/6J CREM^-/-^) and wildtype mice infected by intracecal injection of *E. histolytica*. CREM knockout mice were more susceptible to amebic colitis than wildtype, as assessed by culture of cecal contents (*P*=0.02). **(Figure 4A)** Apoptotic death of cecal intestinal epithelial cells, as detected by caspase 3 stain, was also higher within the CREM^-/-^ mice (*P*=0.028). **(Figures 4B, 4C)**

**Figure 4:**
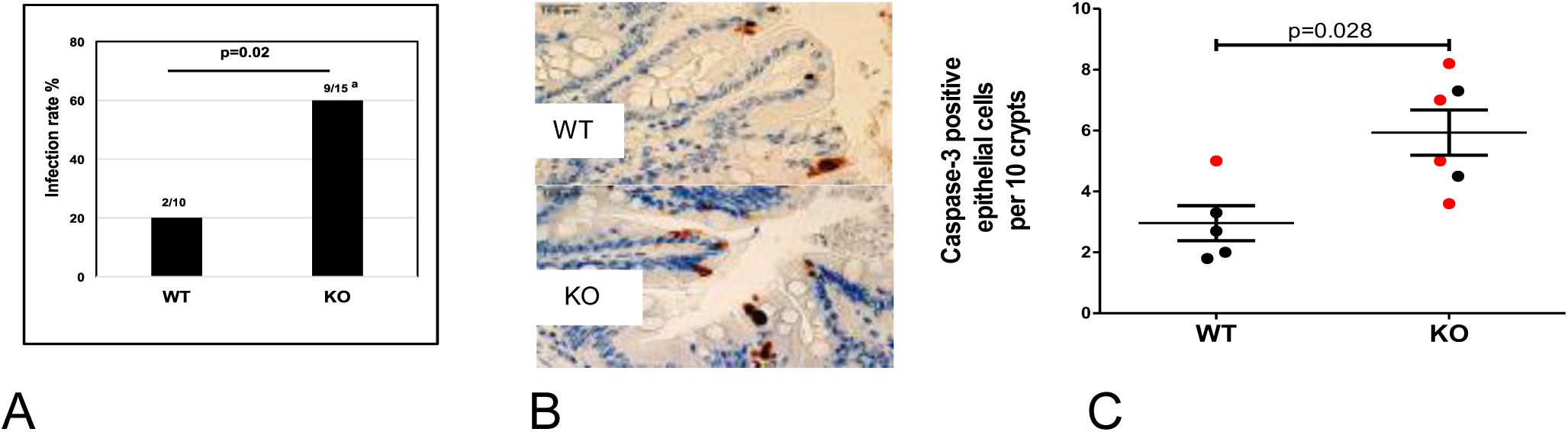
CREM-/-mice (KO) are more susceptible to amebic colitis. **(A)** Mice were infected with *E. histolytica* and euthanized based on clinical score or at 3 days and infection assessed by culture of cecal contents. Numbers above each column represent culture positive samples among total in that group. ^a^ two mice in this group died before 72 h time point. **(b)** Apoptotic death of cecal intestinal epithelial cells was detected by caspase-3 stain and **(C)** scored in a blinded manner (red circles indicate culture positive samples). CREM^-/-^ mice from Blendy et al. lack DNA binding domain 1a-1b and CREM and ICER deficient (Abhyankar & Petri, unpublished).

## Discussion

The most important discovery of this work is that genetic variants in the *CREM*/*CUL2* locus are associated with *Entamoeba histolytica* diarrhea. Many of our most associated SNPs are also reported to be associated with inflammatory bowel disease susceptibility.^20^ The *CREM*/*CUL2* region was identified through the first GWAS conducted on an enteric infectious disease. Haplotype analysis and comparison with 1000 Genomes Project data suggest that this particular association is enhanced in South Asian and European populations. Expression quantitative trait loci (eQTL) analyses identified decreased *CREM* and likely not *CUL2* to be associated with amebiasis. Consistent with eQTL data, *CREM* expression was increased in mice during *E. histolytica* infection and CREM^-/-^ mice had heightened susceptibility to amebic colitis.

The role of CREM as a cAMP-mediated transcriptional regulator promises to add to the understanding of intestinal health by delineating a common mechanism of gut inflammation and repair from infectious (amebiasis) and non-infectious (Crohns and ulcerative colitis) insults. The fact that a polymorphism in *CREM* underlies susceptibility to both indicates that they likely share a common pathway of cAMP-dependent gene regulation, likely from an upset of the homeostatic balance of the gut microbiota with mucosal immunity.

The GWAS meta-analysis was performed on diarrhea-associated *Entamoeba histolytica* infection, or amebiasis, examining two separate birth cohorts: the Dhaka Birth Cohort (DBC) and the Performance of Rotavirus and Oral Polio Vaccines in Developing Countries (PROVIDE) study. These studies gave a unique opportunity to study the genetic susceptibility to enteric infection, with active surveillance capturing the majority of pediatric illness within the first year of life for these children in Dhaka, Bangladesh. The GWAS results identified a significant association (*P*_meta_<5x10^−8^) with a SNP on chromosome 10 within a region encompassing the genes *CUL2* (cullin 2) and *CREM* (cAMP responsive element modulator). Each additional risk allele at this loci conferred a drastic 2.5 times increased odds of *E. histolytica*-associated diarrhea within the first year of life. While there is high linkage disequilibrium within this region leading to an associated block of sites, the only known eQTLs for *CREM* overlapped with our signals while none of the known eQTLs for *CUL2* exhibited association with amebiasis. Functional validation showed a relationship between infection with *E. histolytica* and increased expression of CREM. When applied to a mouse model, CREM^-/-^ mice showed increased susceptibility to amebic colitis when compared to wildtype mice. This evidence reinforces the role of *CREM* in symptomatic *E. histolytica* infection.

The *CREM-CUL2* region has previously been implicated in genome-wide association studies of other traits, most notably of Inflammatory Bowel Disease (IBD), as well as Crohn’s disease (CD) and Ulcerative Colitis (UC). Five of these loci overlap between our meta-analysis and previous studies (rs11010067, rs34779708, rs12261843, rs12242110, and rs17582416). **(Table 4)** Among SNPs with risk allele information, the direction of effect for IBD, Crohn’s disease, or ulcerative colitis was in same as the direction of effect for amebiasis. The effect sizes were stronger for amebiasis when compared to IBD, with rs11010067 conferring 1.14 times the odds of Crohn’s disease within Europeans while conferring 1.88 times the odds of amebiasis within the Bangladeshi cohort, suggesting overlap between IBD and *Entamoeba histolytica* infection in this region. This relationship is also confirmed in the clinical literature. It has previously been observed that the gross findings of amoebic colitis can resemble those seen in inflammatory bowel disease, in which amebic colitis patients can be mistakenly diagnosed as UC or CD. ^7,21^ The acute stage of amebic colitis especially mimics the first attack of colonic Crohn’s disease.^22^ These results suggest that there may be a shared pathway for pathogenesis of infection for amebiasis and autoimmunity for Crohn’s disease that includes *CREM*. The genetic relationship between infection and autoimmune disease it not unprecedented, as leprosy and Crohn’s disease are known to share similar genes also.^23^ The implications of this finding are intriguing especially since the risk allele or haplotype is common in South Asian and European populations.

**Table 3:** Top associations from GWAS meta-analysis.

| rsID | Chr | Position | A0 | A1 | Allele Frequency | OR<sub>DBC</sub> [95%CI] | P<sub>DBC</sub> | OR<sub>PROVIDE</sub> [95%CI] | P<sub>PROVIDE</sub> | OR<sub>META</sub> [95%CI] | P<sub>META</sub> | P<sub>het</sub> |
| --- | --- | --- | --- | --- | --- | --- | --- | --- | --- | --- | --- | --- |
| rs58000832 | 10 | 35517635 | C | CA | 0.227443 | 2.42 [1.54,3.8] | 1.20x10<sup>-4</sup> | 2.56 [1.76,3.73] | 8.50x10<sup>-7</sup> | 2.5 [1.88,3.34] | 4.20x10<sup>-10</sup> | 0.85 |
| rs58994923 | 10 | 35496699 | TA | T | 0.208604 | 2.69 [1.7,4.26] | 2.40x10<sup>-5</sup> | 2.3 [1.58,3.35] | 1.30x10<sup>-5</sup> | 2.45 [1.83,3.27] | 1.50x10<sup>-9</sup> | 0.60 |
| rs11599920 | 10 | 35507164 | G | T | 0.207008 | 2.63 [1.67,4.14] | 3.20x10<sup>-5</sup> | 2.31 [1.59,3.36] | 1.20x10<sup>-5</sup> | 2.43 [1.82,3.25] | 1.70x10<sup>-9</sup> | 0.67 |
| rs75152185 | 10 | 35511869 | G | C | 0.207519 | 2.56 [1.63,4.03] | 4.50x10<sup>-5</sup> | 2.31 [1.59,3.37] | 1.10x10<sup>-5</sup> | 2.41 [1.81,3.22] | 2.10x10<sup>-9</sup> | 0.73 |

**Table 4:** Known GWAS associations in this region show relationship with IBD. (IBD: Inflammatory Bowel Disease, UC: Ulcerative Colitis, CD: Crohn’s Disease)

| SNP | Functional Class | Disease | Allele | P | OR | P<sub>EH</sub> | OR<sub>EH</sub> |
| --- | --- | --- | --- | --- | --- | --- | --- |
| rs11010067 | Downstream gene variant | CD<sup>49</sup> | G | 1x10-26 | 1.14 | 8.7x10-7 | 1.88 |
| rs11010067 | Downstream gene variant | IBD<sup>50</sup> | G | 2x10-25 | 1.12 | 8.7x10-7 | 1.88 |
| rs34779708 | Intron (CREM) | IBD<sup>49</sup> | G | 2x10-25 | NR | 2.1x10-7 | 1.94 |
| rs12261843 | Intron (CCNY) | UC<sup>51</sup> | G | 7x10-7 | 1.07 | 2.6x10-5 | 1.74 |
| rs12242110 | Upstream (CREM) | CD<sup>52</sup> | G | 1x10-9 | 1.15 | 2.5x10-5 | 1.74 |
| rs17582416 | Intergenic | CD<sup>20</sup> | G | 2x10-9 | 1.16 | 1.1x10-6 | 1.87 |

The relationship between the intestinal autoimmune diseases, *E. histolytica*-related diarrhea, and *CREM* may be along the T_h_17 pathway. A transcriptional repressor isoform of *CREM* is denoted *ICER* (inducible cAMP early repressor) and is regulated through a promoter located in one of *CREM*’s introns. Naive *ICER*/*CREM*-deficient CD4+ Þ T-cells have impaired functionality to differentiate to T_h_17, however this can be rescued by forced over-expression of *ICER* specifically.^24^ This relationship is consistent with the role of T_h_17 cells within autoimmune and inflammatory diseases. Specifically, it was found that *ICER*/*CREM*-deficient B6.lpr mice are protected from developing autoimmunity. In addition, high levels of T_h_17 have been found in the gut of subjects with Crohn’s disease.^25^ Mouse models have also shown that intracecal amebic infection resulted in the up regulation of T_h_17 cytokine responses, to the detriment of T_h_1 cytokines.^26^ Further work is needed to elucidate this relationship.

There are several limitations to our analysis. We did not replicate the previously identified association with the leptin receptor, likely due to a different case definition from the original discovery analysis.^11^ This previous study looked at infections (both symptomatic and asymptomatic) within children through the preschool years, while our analysis only examined diarrhea-associated infections within the first year of life. Additional limitations may be overcome in future work, such as suboptimal power due to the small sample sizes of both studies ad our constraints in evaluating the role of co-pathogens. However, because the effect estimates are strong (odds ratio of 2.5 for the most associated risk allele), the combined meta-analysis allowed us to elucidate genome-wide significant associations.

In conclusion, through a meta-analysis of two separate birth cohorts, we uncovered genome-wide significant associations with diarrhea-associated *Entamoeba histolytica* infection within Bangladeshi infants. The top association was on a haplotype spanning *CUL2* and *CREM* on chromosome 10, a region that has previously been implicated with inflammatory bowel disease, Crohn’s disease and ulcerative colitis. Functional evidence suggests a role for *CREM* during early infection with *E. histolytica* and *CREM* knockout mice were more susceptible to amebic colitis. The relationship between infection with the amoebic parasite *Entamoeba histolytica* within the developing world and the development of intestinal autoimmune disorders in the developed world warrants further research to understand their parallels and to expand their respective treatment options.

## Methods

### Dhaka Birth Cohort study design

The Dhaka Birth Cohort (DBC) was part of a longitudinal birth cohort recruited from the urban slum in the Mirpur Thana in Dhaka, Bangladesh. Children were enrolled within the first week after birth, beginning in January 2008, and followed-up bi-weekly with household visits for the first year of life. Trained field research assistants took anthropometric measurements at the time of enrollment, and every three months thereafter. The Height-for-Age Z-score (HAZ) and the Weight-for-Age Z-score (WAZ) were calculated by comparing the height and weight of the study participants with the World Health Organization (WHO) reference population, standardized for age and sex, using the WHO Anthro software, version 3.0.1. Diarrheal stool samples were collected from the home or the study field clinic every time the mother reported diarrhea. These samples were then transported to the Centre for Diarrhoeal Disease Research, Bangladesh (ICDDR,B) parasitology laboratory, maintaining a cold chain. The presence of *E. histolytica* was determined using real-time polymerase chain reaction (RT-PCR), as well as enzyme-linked immunosorbent assay (ELISA). Children were defined as a “case” if they had at least one diarrheal sample that was positive for *E. histolytica* within the first year of life by either method. Children were defined as “controls” if they had no diarrheal samples that were positive for *E. histolytica* within the first year of life by either method, and also had at least one diarrheal or monthly stool sample tested for a true negative control.

### Dhaka Birth Cohort genotyping

The samples from DBC were genotyped as part of a larger set of 1,573 samples from four Bangladesh study groups. These samples were genotyped in three separate batches at the University of Virginia Center for Public Health Genomics Laboratory. Sample preparation and genotype calling followed standard Illumina protocols. Only the 484 DBC samples genotyped across three batches were used for this analysis: 165 were genotyped in batch 1 (Illumina Human1M-duoV3; 1,199,187 total SNPs/CNV sites); 154 in batch 2 (Illumina HumanOmni1-Quad v1.0: 1,140,419 total SNPs/CNV probes); and 165 in batch 3 (Illumina HumanOmni2.5-4v1: 2,450,000 total SNPs/CNV probes). Samples were dropped from the analysis if: 1) genotyping call rate was < 95%; 2) they were cryptic duplicates with differing phenotype records or were cryptically related up to first degree in the same study group, relationships inferred with KING^27^; 3) the inferred sex from the genetic X/Y chromosome data did not match study database gender. SNPs were dropped if: 1) per batch call rate was < 95%; 2) per batch p-value for test of Hardy-Weinberg proportions < 1 x 10 −4 (X chr females only); 3) they were identified as CNV probes; 4) SNPs mapped to multiple locations in the genome. We remapped all SNPs to Human Genome Build 37 and merged individual batches of data into a single combined data set with 529,893 common intersecting SNPs in the 3 batches by rsID. The full data set was imputed to the 1000 Genomes Phase 3 reference data^16^ with phasing through SHAPEIT^28,29^ and imputation with IMPUTE2^30-34^.

### PROVIDE study design

The “Performance of Rotavirus and Oral Polio Vaccines in Developing Countries” (PROVIDE) Study is a randomized controlled clinical trial birth cohort that was designed to evaluate factors that may influence oral vaccine efficacy among children from areas with high poverty, urban overcrowding, and poor sanitation.^15^ A total of 700 children and their mothers were followed for the child’s first 2 years of life with a 2x2 factorial design looking specifically at the efficacy of 2-dose Rotarix oral rotavirus vaccine and oral polio vaccine (OPV) with an inactivated polio vaccine (IPV) boost. This study was performed with the International Center for Diarrheal Disease Research, Bangladesh (ICDDR,B) with the study population all from the Mirpur area of Dhaka, Bangladesh. Pregnant mothers were recruited from the community by female Bangladeshi field research assistants (FRAs). Participants had fifteen scheduled follow-up clinic visits, with biweekly diarrhea surveillance at their homes by FRAs. The presence of *E. histolytica* in the diarrheal samples was determined by RT-PCR. Children were defined as a “case" if they had at least one diarrheal sample that was positive for *E. histolytica* within the first year of life. Children were defined as “controls” if they had at least one diarrheal sample available for testing, but none were positive for *E. histolytica*.

### PROVIDE genetic data

Within PROVIDE, 541 children were genotyped on Illumina’s Infinium Multiethnic Global Array (MEGA). Standard quality control metrics were used for the genome-wide data. Single nucleotide polymorphism (SNP) filters included genotype missingness <5% (none), minor allele frequency (MAF) >0.5% (M=659,171), and Hardy-Weinberg equilibrium P-value >10E-5 (M=789). Individuals were filtered for individual missingness <2% (none), heterozygosity outliers (N=4), principal components outliers (none). One individual from each first and second degree relative pairs were removed (N=36). After both individual and SNP-level filters, there were 699,246 SNPs and 499 individuals. The genetic data was split into chromosomes for phasing and imputation. Each chromosome was phased using SHAPEIT^28,29^ v2.r790 with 1000 Genomes Project Phase 3 data as the reference.^16^ After phasing, the chromosomes were imputed using IMPUTE v2.3.2^30-34^ using 1000 Genomes Project Phase 3 data as reference.

### *E. histolytica* detection protocol

The detection protocol for E. histolytica has previously been described in Haque et al (2007). ^35^ Primers and Taqman probes for E. histolytica (accession no. X64142) were designed on a small subunit ribosomal RNA gene, with the amplified targets at 134, 62, and 151 bp. All primers and Taqman probes were purchased from Eurogentec (Seraing, Belgium). Multiplex real-time PCR was conducted using standard protocol previously described.

### Association analyses

To estimate the associations between genetics and diarrhea-associated E. *histolytica* infection, each study (DBC and PROVIDE) was initially run separately. An unadjusted logistic regression was run with SNPTEST^30,34,36^, incorporating the imputed genotypes’ weights and assuming an additive model of inheritance. The two studies were incorporated into a meta-analysis using the software META^37^ in a fixed-effects analysis. Results were then filtered for MAF > 5%, INFO > 60% in both cohorts, and a P-value for heterogeneity between the two cohorts greater than 0.01. This ensured stable estimates of association that were adequately powered in both analyses separately and together.

### eQTL analysis

Known expression quantitative trait loci (eQTLs) from the Genotype-Tissue Expression project (GTEx) portal (www.gtexportal.org) were used.^17^ We used the association results from the GTEx eQTL analyses for all tissues for *CREM*, *CUL2*, and *CCNY*. The overlap between these association results and the imputed meta-analysis results were included (*CREM* (EBV-transformed lymphocytes): 490 sites, *CUL2* (Whole Blood Cells-Transformed fibroblasts): 2 sites, *CCNY* (Esophagus-Mucosa): 80 sites).

### Haplotype analyses

For all haplotype analyses, only the genotyped original PROVIDE data was used to avoid potential bias resulting from imputation. The region of association on chromosome 10 between 35.25mb and 35.55mb was subset from the previously phased data, including both *CREM* and *CUL2*. A total of 26 SNPs were included in this one large haplotype block. A chi-squared test was used to compare the distribution of outcome in individuals who had a copy of the index haplotype versus those without any copy of the index haplotype. The top associations were then assessed using logistic regression to estimate effect size on an individual, instead of haplotype. The top associated haplotype was then compared against reference populations within the 1000 Genomes Project (TGP).^16^ For each superpopulation (Africa, Americas, East Asia, Europe, and South Asia), the 26 SNPs were subset from previously phased data. There were 23 unique haplotypes found within TGP that occurred at least 10 times (on 10 separate chromosomes). The haplotype network was mapped within R, using an infinite site model of DNA sequences from the pairwise deletion of missing data within the library “pegas”. Enrichment for a haplotype at a continental level was assessed using a chi-squared test.

### Selection analyses

The 1000 Genomes Project (TGP) data for chromosome 10 was assessed for evidence of selection using four representative populations: Bengalis in Bangladesh, Western Europeans in the United States, Han Chinese in Beijing China, and Yoruba in Nigeria. These four populations were selected to be representative of the different continents, as well as the source population for both DBC and PROVIDE. To determine the presence of positive selection, the integrated haplotype score (iHS) was calculated using selscan. ^38-42^ To assess the presence of balancing or directional selection, Tajima’s D was calculated within vcftools.^43^

### *E. histolytica* culture

*E. histolytica* trophozoites were maintained in a trypsin-yeast extract-iron (TYI-S-33) medium supplemented with 2% Diamond vitamins, 13% heat inactivated bovine serum (Gemini Labs) and 100 U/ml penicillin plus 100 μg/ml streptomycin (Invitrogen). ^44^ Trophozoites originally derived from HM1: IMSS (ATCC) and passed sequentially through mice to maintain animal virulence were used for challenge experiments.

### CRE-Luciferase reporter assays

HCT116 cells were maintained in McCoy’s media supplemented with 10% heat-inactivated fetal bovine serum. Cells were transfected with pCRE Tluc16-DD Vector (Thermo Fisher Scientific (88247) using lipofectamine 2000 (Thermo Fisher Scientific 11668019). The pCRE Tluc16-DD vector contains an optimized minimal core promoter and 5 tandem repeats of the cAMP response element (CRE) a turboluciferase reporter with a dual-destabilization domain. Luciferase levels were measured using the TurboLuc™ Luciferase One-Step Glow Assay Kit (Thermo 88263). Timecourse assays were done with cells exposed to *E. histolytica* at a ratio of 1 parasite to 5 host cells in transwells to delay contact-dependent cytotoxicity. CREM luciferase assays were done in cells co-transfected with was esiRNA EHU125161 targeting the 2nd DNA binding domain CREM (Sigma) or GFP as a control. *E. histolytica* secreted products were obtained as previously described.^45^ % induction was calculated (mean of biological replicates from experimental condition/mean of biological replicates of the media control) * 100. Error bars representing standard error are shown. P-values were calculated by unpaired t-test with no correction for multiple comparisons using Prism 6.0 (Graphpad).

### Q223R mice

All microarray data discussed in this paper were deposited into NCBI’s Gene Expression Omnibus ^46^ and are accessible through GEO Series accession number GSE43372 (Mackey-lawerence et al). P-values were calculated by unpaired t-test with no correction for multiple comparisons using Prism 6.0 (Graphpad).

### Challenge experiments using ICER/ CREM^-/-^ mice

C57BL/6J.CREM^-/-^mice were derived from SV129/Bl6.CREM^-/-^ mice in which the CREM DNA binding domains were replaced by a LacZ-neo fusion cassette, as originally cloned by Blendy et al. ^47^ These mice were crossed to C57BL/6J mice for over nine generations (Yoshida, et.al. 2016). All experiments were done strictly according to the IACUC, University of Virginia guidelines and using approved protocol. Infection with *E. histolytica* was performed on C57BL/6J.ICER/CREM^-/-^ mice and littermate wild type controls. Both male and female mice, between 7-18 weeks of age were used. Mice were regenotyped to validate. Since male mice homozygous for CREM^-/-^ are sterile, we maintained a heterozygous breeding colony. Trophozoites originally derived from HM1:IMSS (ATCC) and passed sequentially through mice to maintain animal virulence were used for infection. Mice received a cocktail of four antibiotics (1g/L each ampicillin, neomycin and metronidazole; 0.5g/L vancomycin) in drinking water for two weeks prior to infection. Metronidazole was omitted from the cocktail four days prior to challenge. Mice were challenged intracecally with two million trophozoites in 150 μl medium following laparotomy. ^48^ Mice were euthanized based on their clinical score or on day-3 post challenge, whichever came first. Cecal contents were suspended in 1 ml PBS and 300 μl used for culturing in TYI-S-33 broth for up to five days. Infection rates were analyzed using Chi-square test.

### Caspase-3 immunostaining

Mouse cecal tissue was fixed in Bouin’s solution for 24h and washed with 70% ethanol. Paraffin embedded cecal sections were stained at the biorepository core facility of the University of Virginia using cleaved caspase-3 specific antibody (Cell Signaling, Cat. # 9661L). Number of caspase-3 positive (brown) epithelial cells and crypts were scored in a blinded fashion. Data was analyzed using Mann-Whitney test.

## Acknowledgements

This work was funded by grants to WP from the Bill & Melinda Gates Foundation and the National Institutes of Health, Allergy and Infectious Disease AI043596 and to Priya Duggal from the Sherrilyn and Ken Fisher Center for Environmental Infectious Diseases Discovery Program. The funders had no role in the study design, data collection and data analysis, decision to publish, or preparation of the manuscript. Icddr,b is grateful to the governments of Bangladesh, Canada, Sweden, and the UK for providing core unrestricted support. We thank the families of the Mirpur field area who participated in this study, and we also thank the world of the field and lab staffs of the Parisitology Laboratory of icddr,b who worked for the Dhaka Birth Cohort (DBC) and PROVIDE projects, without whom we could not have completed this research.

## Bibliography

1. United Nations. The Millennium Development Goals Report. 1–15 (2015).

2. Liu, L. et al. Global, regional, and national causes of child mortality in 2000–13, with projections to inform post-2015 priorities: an updated systematic analysis. The Lancet (2015). doi:10.1016/S0140-6736(14)61698-6

3. Haque, R. et al. Entamoeba histolytica Infection in Children and Protection from Subsequent Amebiasis. Infection and Immunity 74, 904–909 (2006).

4. Mondal, D., Haque, R., Sack, R. B., Kirkpatrick, B. D. & Petri, W. A. Attribution of malnutrition to cause-specific diarrheal illness: evidence from a prospective study of preschool children in Mirpur, Dhaka, Bangladesh. Am. J. Trop. Med. Hyg. 80, 824–826 (2009).

5. Mondal, D., Petri, W. A., Sack, R. B., Kirkpatrick, B. D. & Haque, R. Entamoeba histolytica-associated diarrheal illness is negatively associated with the growth of preschool children: evidence from a prospective study. Trans. R. Soc. Trop. Med. Hyg. 100, 1032–1038 (2006).

6. Haque, R., Huston, C. D., Hughes, M., Houpt, E. & Petri, W. A. Amebiasis. N. Engl. J. Med. 348, 1565–1573 (2003).

7. Stanley, S. L. Amoebiasis. The Lancet 361, 1025–1034 (2003).

8. Marie, C. & Petri, W. A. Regulation of virulence of Entamoeba histolytica. Annu. Rev. Microbiol. 68, 493–520 (2014).

9. Mondal, D. et al. Contribution of Enteric Infection, Altered Intestinal Barrier Function, and Maternal Malnutrition to Infant Malnutrition in Bangladesh. Clinical Infectious Diseases 54, 185–192 (2011).

10. Watanabe, K. & Petri, W. A. Molecular biology research to benefit patients with Entamoeba histolytica infection. Mol. Microbiol. 98, 208–217 (2015).

11. Duggal, P. et al. A mutation in the leptin receptor is associated with Entamoeba histolytica infection in children. J. Clin. Invest. 121, 1191–1198 (2011).

12. Mackey-Lawrence, N. M. et al. Effect of the leptin receptor Q223R polymorphism on the host transcriptome following infection with Entamoeba histolytica. Infection and Immunity 81, 1460–1470 (2013).

13. Marie, C. S., Verkerke, H. P., Paul, S. N., Mackey, A. J. & Petri, W. A. Leptin protects host cells from Entamoeba histolytica cytotoxicity by a STAT3-dependent mechanism. Infection and Immunity 80, 1934–1943 (2012).

14. Duggal, P. et al. Influence of human leukocyte antigen class II alleles on susceptibility to Entamoeba histolytica infection in Bangladeshi children. J. Infect. Dis. 189, 520–526 (2004).

15. Kirkpatrick, B. D. et al. The ’Performance of Rotavirus and Oral Polio Vaccines in Developing Countries’ (PROVIDE) Study: Description of Methods of an Interventional Study Designed to Explore Complex Biologic Problems. American Journal of Tropical Medicine and Hygiene 1–17 (2015). doi:10.4269/ajtmh.14–0518

16. Auton, A. et al. A global reference for human genetic variation. Nature 526, 68–74 (2015).

17. GTEx Consortium & Kellis, M. Human genomics. The Genotype-Tissue Expression (GTEx) pilot analysis: multitissue gene regulation in humans. Science 348, 648–660 (2015).

18. McGowan, K., Piver, G., Stoff, J. S. & Donowitz, M. Role of prostaglandins and calcium in the effects of Entamoeba histolytica on colonic electrolyte transport. YGAST 98, 873–880 (1990).

19. Rico, G., Diaz-Guerra, O. & Kretschmer, R. R. Cyclic nucleotide changes induced in human leukocytes by a product of axenically grown Entamoeba histolytica that inhibits human monocyte locomotion. Parasitol Res 81, 158–162 (1995).

20. Barrett, J. C. et al. Genome-wide association defines more than 30 distinct susceptibility loci for Crohn’s disease. Nat Genet 40, 955–962 (2008).

21. Tucker, P. C., Webster, P. D. & Kilpatrick, Z. M. Amebic colitis mistaken for inflammatory bowel disease. Arch. Intern. Med. 135, 681–685 (1975).

22. De Hertogh, G. & Geboes, K. Crohn’s disease and infections: a complex relationship. MedGenMed 6, 14 (2004).

23. Zhang, F.-R. et al. Genomewide Association Study of Leprosy. N. Engl. J. Med. 361, 2609– 2618 (2009).

24. Yoshida, N. et al. ICER is requisite for Th17 differentiation. Nature Communications 7, 12993 (2016).

25. Annunziato, F. et al. Phenotypic and functional features of human Th17 cells. Journal of Experimental Medicine 204, 1849–1861 (2007).

26. Guo, X., Stroup, S. E. & Houpt, E. R. Persistence of Entamoeba histolytica infection in CBA mice owes to intestinal IL-4 production and inhibition of protective IFN-?. Mucosal Immunol 1, 139–146 (2008).

27. Manichaikul, A. et al. Robust relationship inference in genome-wide association studies. Bioinformatics 26, 2867–2873 (2010).

28. Delaneau, O., Marchini, J. 1000 Genomes Project Consortium. Integrating sequence and array data to create an improved 1000 Genomes Project haplotype reference panel. Nature Communications 5, 3934 (2014).

29. Delaneau, O., Zagury, J.-F. & Marchini, J. correspondence. Nature Methods 10, 5–6 (2013).

30. Marchini, J., Howie, B., Myers, S., McVean, G. & Donnelly, P. A new multipoint method for genome-wide association studies by imputation of genotypes. Nature Genetics 39, 906–913 (2007).

31. Howie, B., Fuchsberger, C., Stephens, M., Marchini, J. & Abecasis, G. R. Fast and accurate genotype imputation in genome-wide association studies through pre-phasing. Nat Genet 44, 955–959 (2012).

32. Howie, B., Marchini, J. & Stephens, M. Genotype imputation with thousands of genomes. G3: Genes (2011). doi:10.1534/g3.111.001198/-/DC1

33. Howie, B. N., Donnelly, P. & Marchini, J. A flexible and accurate genotype imputation method for the next generation of genome-wide association studies. PLoS Genetics 5, e1000529 (2009).

34. Marchini, J. & Howie, B. Genotype imputation for genome-wide association studies. Nature Reviews Genetics 11, 499–511 (2010).

35. Haque, R. et al. Multiplex real-time PCR assay for detection of Entamoeba histolytica, Giardia intestinalis, and Cryptosporidium spp. American Journal of Tropical Medicine and Hygiene 76, 713–717 (2007).

36. Burton, P. R. et al. Genome-wide association study of 14,000 cases of seven common diseases and 3,000 shared controls. Nature 447, 661–678 (2007).

37. Liu, J. Z. et al. Meta-analysis and imputation refines the association of 15q25 with smoking quantity. Nature Genetics 42, 436–440 (2010).

38. Sabeti, P. C. et al. Genome-wide detection and characterization of positive selection in human populations. Nature 449, 913–918 (2007).

39. Ferrer-Admetlla, A., Liang, M., Korneliussen, T. & Nielsen, R. On detecting incomplete soft or hard selective sweeps using haplotype structure. Molecular Biology and Evolution 31, 1275–1291 (2014).

40. Szpiech, Z. A. & Hernandez, R. D. selscan: an efficient multithreaded program to perform EHH-based scans for positive selection. Molecular Biology and Evolution 31, 2824–2827 (2014).

41. Sabeti, P. C. et al. Detecting recent positive selection in the human genome from haplotype structure. Nature 419, 832–837 (2002).

42. Voight, B. F., Kudaravalli, S., Wen, X. & Pritchard, J. K. A Map of Recent Positive Selection in the Human Genome. PLoS Biol 4, e72 (2006).

43. Danecek, P. et al. The variant call format and VCFtools. Bioinformatics 27, 2156–2158 (2011).

44. Diamond, L. S., Harlow, D. R. & Cunnick, C. C. A new medium for the axenic cultivation of Entamoeba histolytica and other Entamoeba. Trans. R. Soc. Trop. Med. Hyg. 72, 431–432 (1978).

45. Yu, Y. & Chadee, K. Entamoeba histolytica stimulates interleukin 8 from human colonic epithelial cells without parasite-enterocyte contact. YGAST 112, 1536–1547 (1997).

46. Edgar, R., Domrachev, M. & Lash, A. E. Gene Expression Omnibus: NCBI gene expression and hybridization array data repository. Nucleic Acids Research 30, 207–210 (2002).

47. Blendy, J. A., Kaestner, K. H., Weinbauer, G. F., Nieschlag, E. & Schutz, G. Severe impairment of spermatogenesis in mice lacking the CREM gene. Nature 380, 162–165 (1996).

48. Houpt, E. et al. Prevention of intestinal amebiasis by vaccination with the Entamoeba histolytica Gal/GalNac lectin. Vaccine 22, 611–617 (2004).

49. Liu, J. Z. et al. Association analyses identify 38 susceptibility loci for inflammatory bowel disease and highlight shared genetic risk across populations. Nat Genet (2015). doi:10.1038/ng.3359

50. Jostins, L. et al. Host-microbe interactions have shaped the genetic architecture of inflammatory bowel disease. Nature 491, 119–124 (2012).

51. Anderson, C. A. et al. Meta-analysis identifies 29 additional ulcerative colitis risk loci, increasing the number of confirmed associations to 47. Nat Genet 43, 246–252 (2011).

52. Franke, A. et al. Genome-wide meta-analysis increases to 71 the number of confirmed Crohn’s disease susceptibility loci. Nature Publishing Group 42, 1118–1125 (2010).

